# G-Quadruplexes Act as an On/Off Switch While i-Motifs Regulate Insulin Expression in Reporter Gene Assays

**DOI:** 10.1101/2025.05.02.651924

**Authors:** Dilek Guneri, Christopher J. Morris, Yiliang Ding, Timothy D. Craggs, Steven S. Smith, Zoë A. E. Waller

## Abstract

The insulin-linked polymorphic region (ILPR) is a variable number tandem repeat located in the promoter of the human insulin gene. This G-rich sequence can fold into four-stranded G-quadruplex DNA structures, while its complementary C-rich strand forms i-motifs. The ILPR varies in repeat number and sequence composition, but the relationship between sequence diversity, DNA structure, and insulin gene regulation remains poorly understood. Although both G-quadruplexes and i-motifs have been implicated in transcriptional control, their relative contributions, particularly when formed on complementary strands of the same locus, are unclear. Here, we characterised the structure and stability of nine ILPR-based sequences using biophysical techniques and luciferase reporter assays. We demonstrate that transcriptional activation in response to high glucose occurs only when both G-quadruplex and i-motif structures can form. Other combinations of structures do not induce transcription. Moreover, promoter activity correlated positively with i-motif stability, but not with G-quadruplex stability. These results suggest a model in which G-quadruplexes function as an on/off switch, while i-motifs act as modulators of gene expression. Our findings underscore the importance of treating G-quadruplexes and i-motifs as a dynamic, interdependent system in both the regulation of gene expression and also the potential of these structures as therapeutic targets.

**Graphical Abstract:** 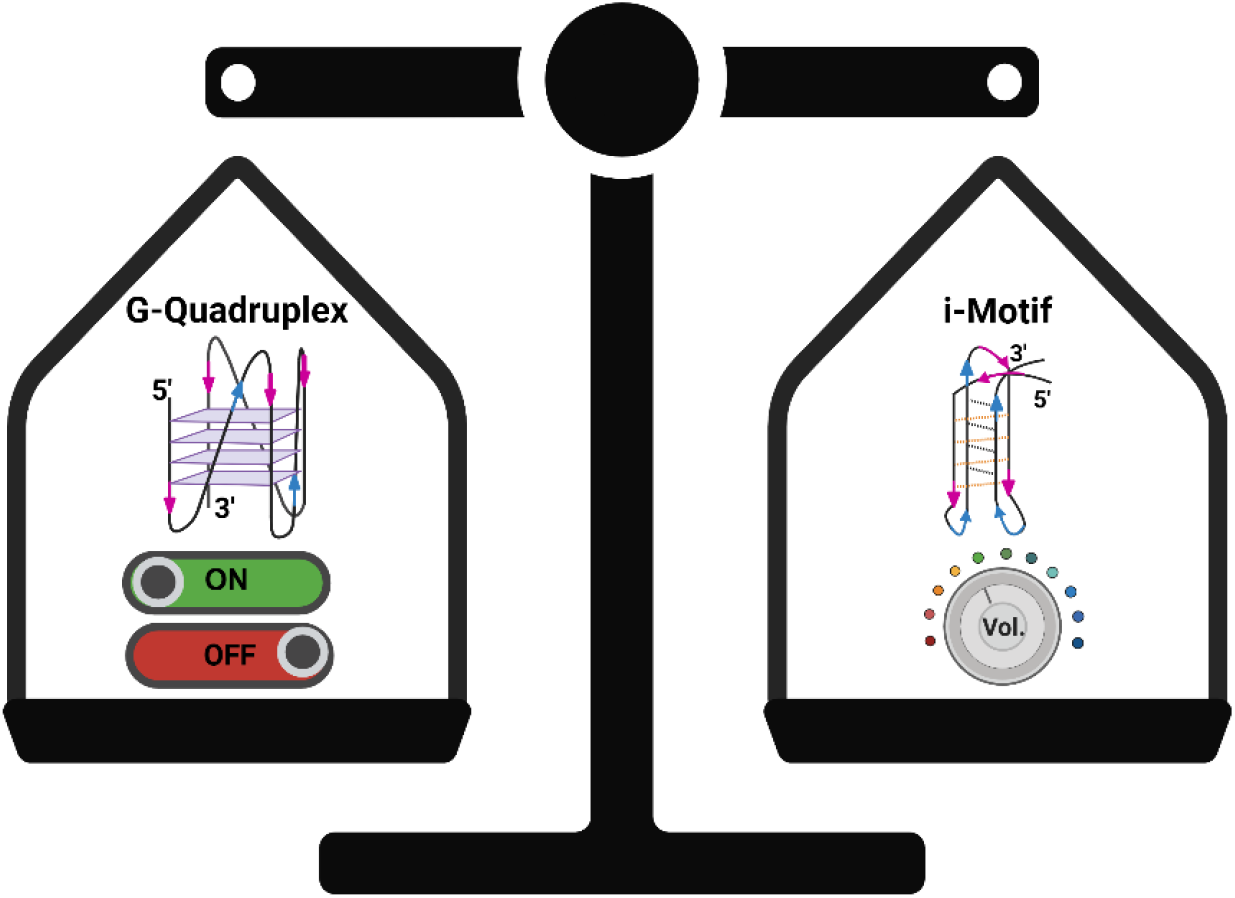

## Introduction

Insulin is a central protein hormone regulating glucose metabolism. Deficiencies of insulin production can cause hyperglycaemia and lead to diabetes mellitus [1, 2]. The insulin-linked polymorphic region (ILPR) is found 363 bp upstream of the human insulin transcription start site [3-5]. This heterogeneous region may vary in nucleotide composition and the number of tandemly repeated ILPR sequences among individuals [2, 5-8]. A shortened ILPR repeat length is linked with Type-1 Diabetes [1] while some variations in the sequence composition is linked to Type-2 Diabetes [2, 5, 7-10]. The ILPR is found to regulate insulin as well as insulinlike growth factor 2 and small variations in the ILPR sequence are associated with decreased gene expression in both genes [7, 11, 12].

The ILPR is comprised of a variable number of tandem repeats with sequence variability, the predominant sequence being 5’-ACAGGGGTGTGGGG-3’/3’-TGTCCCCACACCCC-5’ [5, 7, 8], This GC-rich DNA region has the potential to form non-canonical DNA structures. The Crich sequence can fold into i-motif structures that are stabilised by pH, negative supercoiling, and molecular crowding [13-17]. The complementary G-rich sequence is arranged as planar G-quartets held together by Hoogsteen hydrogen-bonding and stabilised via π–π stacking and cations [18-20]. These types of non-canonical DNA structures have been shown, in cells, to be prevalent in the promotor regions of genes and to appear at certain cellular events [5, 21-24]. Recently, we characterised 11 of the most common native ILPR variants and established a relationship between i-motif and G-quadruplex structure formation regulating in celluo reporter gene expression [25]. This then led to the question of which is more important for insulin expression: G-quadruplex or i-motif structure?

Early research showed that ILPR variants influence Pur-1–mediated gene expression, with the predominant sequence eliciting the strongest response, speculatively due to the formation of inter- and intramolecular G-quadruplex structures [8]. Chemical foot-printing and mechanical folding experiments have indicated that both G-quadruplex and i-motif structures can form within the double-stranded ILPR region, but the structures are mutually exclusive [26]. However, more recently, we were able to demonstrate a link between the non-canonical DNA structure formation and regulation of gene expression [25]. Based on the biophysical data and biological experiments, we sought to determine which structure plays a more important role in regulating insulin reporter expression.

Herein, we determine a precise relationship between non-canonical DNA structure formation and stability, and the expression of the ILPR-regulated reporter gene. Through a systematic study of mutant ILPR sequences we show that both G-quadruplex and i-motif are essential for insulin reporter expression and the level of expression correlates positively with i-motif, but not G-quadruplex, stability.

## Materials and Methods

### Oligonucleotides

All synthetic oligonucleotide sequences used in this study were synthesised and reverse-phase HPLC purified by Eurogentec (Belgium). Samples were prepared at a final concentration of 1 mM in ultra-pure water and concentrations were confirmed using a Nanodrop spectrophotometer. For DNA structure annealing, samples in buffer, as specified in experimental sections, were heated at 95°C for 5 minutes in a heating block, followed by slow cooling to room temperature overnight.

### Circular Dichroism Spectroscopy

Circular dichroism (CD) spectra of the selected ILPR sequences were recorded using a JASCO J-1500 spectropolarimeter under a constant flow of nitrogen. C-rich ILPR samples were diluted to 10 μM in 10 mM sodium cacodylate buffer (NaCaco, Merck) containing 100 mM KCl, and measured across a pH range of 4 to 8. G-rich ILPR samples were prepared at 10 μM in 10 mM NaCaco with either 100 mM KCl, 100 mM NaCl, or 100 mM LiCl at pH 7.0 to assess cation-specific folding. For each buffer and sample, four spectra scans were accumulated over a range of 200–320 nm at 20 °C, using a data pitch of 0.5 nm, scanning speed of 200 nm/min, 1-second response time, 1 nm bandwidth, and a sensitivity setting of 200 mdeg. Spectra were baseline-corrected at 320 nm. The transitional pH (pH_T_) of the C-rich ILPR sequences was determined from triplicate experiments by fitting a sigmoidal curve to the ellipticity at 288 nm across the pH range, with the inflection point representing the pH_T_.

### UV Melting/Annealing and Thermal Difference Spectroscopy

UV melting/annealing and TDS experiments were carried out using a Jasco V-750 UV–Vis spectrophotometer. C-rich ILPR samples were prepared at 2.5 μM in 10 mM sodium cacodylate (NaCaco) buffer with 100 mM KCl at pH 5.5, while G-rich ILPR samples were annealed at 2.5 μM in 10 mM NaCaco with 20 mM KCl at pH 7.0. Samples were initially held at 4 °C for 10 minutes, then subjected to three thermal cycles of melting and annealing. During melting, the temperature was gradually increased from 4 °C to 95 °C at a rate of 0.5 °C/min, with absorbance recorded at 260 nm and 295 nm in 1 °C intervals, following a 5-minute hold at each temperature. After a 10-minute hold at 95 °C, the process was reversed for annealing.

Absorbance values were baseline-corrected, and the fraction folded was calculated for both wavelengths. The melting temperature (*T*_m_) and annealing temperature (*T*_a_) were determined using the first derivative of the corrected and normalised data for each cycle [27].

For TDS analysis, spectra were recorded between 230 nm and 320 nm after maintaining the samples at 4 °C (folded) and 95 °C (unfolded) for 10 minutes each. The TDS profile was generated by subtracting the folded spectrum from the unfolded spectrum, zero-corrected at 320 nm, and normalised to the maximum absorbance to reveal structure-specific signatures.

### Cell Culture

INS-1 rat insulinoma cells (AddexBio, Catalogue No. C0018007) were cultured in RPMI-1640 medium supplemented with 10% fetal bovine serum (FBS), 50 μM 2-mercaptoethanol (BME), and 1% penicillin-streptomycin (all reagents from Gibco). Cells were maintained at 37 °C in a humidified incubator with 5% CO_2_. The culture medium was replaced every three days, and cells were passaged upon reaching ∼80% confluency. All experiments were performed using cells between passages 10 and 14.

### Transfection of Reporter Gene Plasmids

The firefly luciferase reporter gene was placed under the control of the human insulin promoter containing one of the selected ILPR variants or corresponding mutants. INS-1 cells were cotransfected with the pRL-TK Renilla reference vector under previously described conditions [25]. For data analysis, we adapted the approach described by Baker and Boyce to calculate the promoter induction ratio [28]. Specifically, each Firefly:Renilla ratio was normalised by dividing it by the average Firefly:Renilla ratio of the corresponding plate, excluding the top and bottom 25% of wells to minimise the influence of outliers. This refinement enabled a more accurate distinction between the transcriptional activities of different ILPR variants and mutants in response to either 2.8 mM or 16.2 mM glucose stimulation.

## Results and Discussion

### Selection of ILPR variant sequences for analysis

We designed 36 systematic mutations within the cytosine tract or the loop region of the C-rich ILPR variants, generating a biophysical landscape for a wide range of i-motif stabilities [25, 29]. From this library, we selected nine ILPR sequences, comprising seven naturally-occurring native variants and two systematically modified mutants. For clarity, we refer to the naturally derived sequences as ILPR variants, and to the artificially introduced sequence modifications as ILPR mutants. The transitional pH (pH_T_) was determined from CD spectra recorded across a range of pH values and represents the pH at which 50% of the i-motif population is folded/unfolded. This value serves as a quantitative indicator of i-motif stability [13, 16, 30, 31]. We aimed for a broad range of transitional pHs to enable determination of a potential cut-off point in i-motif stability that may affect gene expression (Figure 1). The selected G/C-rich ILPR sequences are summarised in Table S1 (C-rich variants) and Table S2 (G-rich variants).

**Figure 1.**
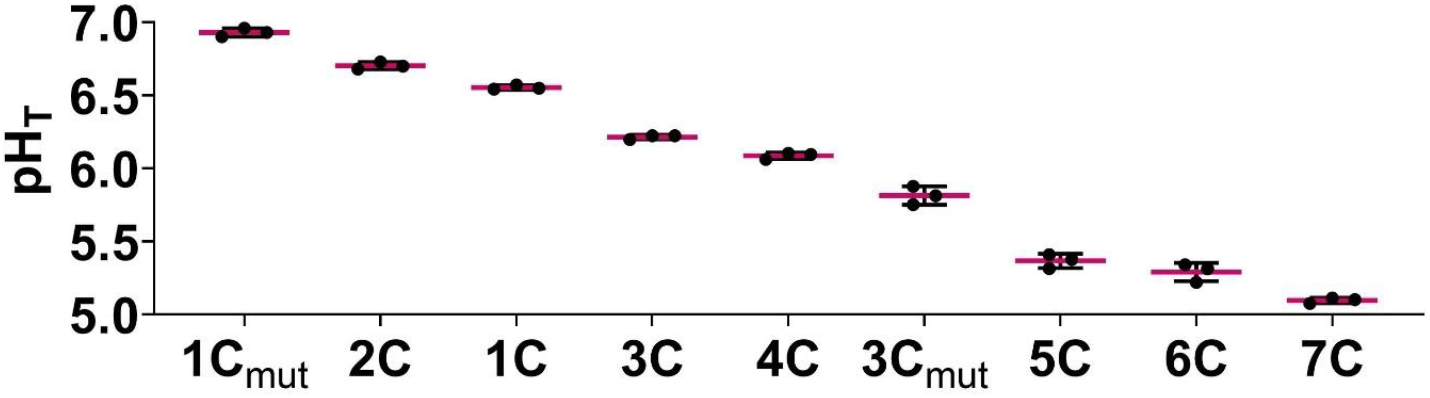
Highest to lowest transitional pH (pH_T_) of selected C-rich ILPR variants including mutants as determined by CD spectroscopy. Data shown as Mean ± SD (n=3). Source data is available.

### Effects of non-native mutations on ILPR variant sequence topologies

Mutations in the loop regions of the prevalent ILPR sequence 1C (from an ACA loop to a TCT) was found to further stabilise the i-motif from pH_T_ 6.6 to pH_T_ 6.9 (Table 1, Figure S1A/F and S1B/G, p <0.0001). This is consistent with other studies that indicated that having TT base pairs in the loops provides additional stability [25][32].

**Table 1.**
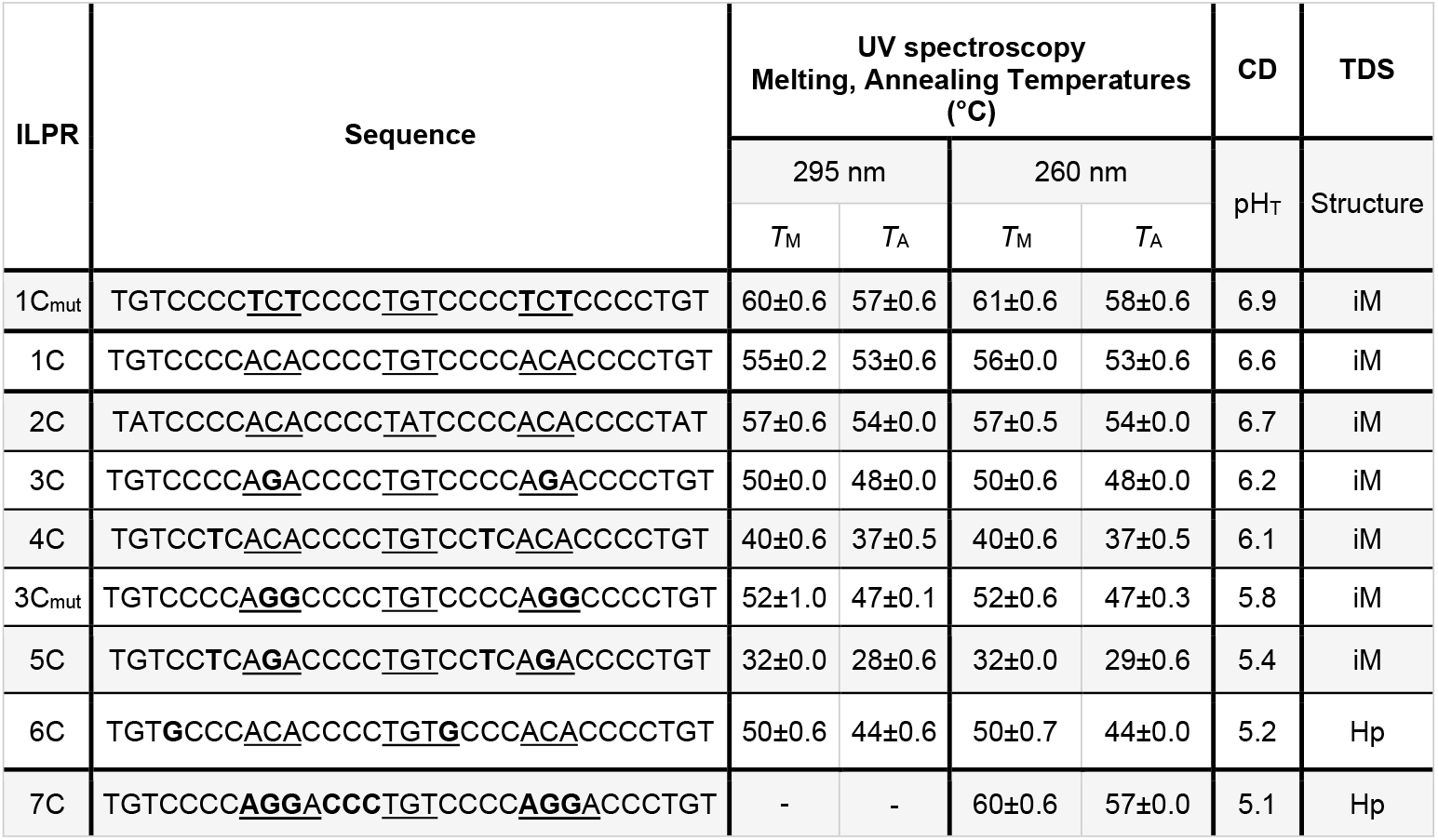
Biophysical characterisation via CD spectrum, UV melt and anneal, and thermal difference spectrum of C-rich ILPR mutants in comparison with naturally occurring ILPR variants as published in Guneri et al., 2024. (− = not detected, T_M_ = Melting Temperature, T_A_= Annealing Temperature, pHT = Transitional pH, i-Motif = iM, Hairpin = Hp)

For example, a DNA microarray of 10,976 potential i-motif–forming sequences revealed that i-motifs with short loops (1–4 nucleotides) were more stable when thymine residues flanked the cytosine tracts [32]. We have also observed TT base pairs in the ILPR structure through the crystal structure, NMR experiments and supporting molecular dynamics simulations of the ILPR i-motif [25]. TT base pairs have previously been shown to stabilise non-canonical DNA structures by forming reverse Hoogsteen hydrogen bonds or loop-stabilising interactions, particularly under mildly acidic or dehydrated conditions, contributing to structural integrity and pH responsiveness in i-motifs and triplexes [33]. The mutation in the loop of ILPR 3C (from AGA to AGG) decreased the transitional pH from 6.2 to 5.8 (Table 1, Figure S1D/I and S1E/J, p <0.0001). This is consistent with findings by Benabou *et al*., which indicated that the incorporation of guanine bases into the loops of an i-motif decrease i-motif stability while thymine or cytosine residues in these positions enhance i-motif stability [34]. Seven out of the nine selected C-rich ILPR variants were found to fold into i-motifs with pH_T_-values between pH 5.4 and 6.9 and these unfold into a what appears to be a random coil. CD spectroscopy of ILPR 1, ILPR 1C_mut_, ILPR 2C, ILPR 3C_mut_, ILPR 4C, and ILPR 5C showed i-motif formation at acidic pH, indicated by a positive peak at 288 nm and a negative peak at 260 nm, and as the pH increases the structure unfolds, shifting the positive peak between 275 - 280 nm and the negative peak to 240 nm (Figure S1 and S2). Typical i-motif spectra have been shown to shift their positive bands from about 288 nm to 280 nm [35-37]. However, ILPR 6C and ILPR 7C have a positive peak at 285 nm which shifts to about 283 nm (Figure 1, Table 1, Figure S2C/G and S2D/H), indicating the formation of hairpins [38, 39]. UV melting/annealing experiments are widely used to determine the thermal stability [27, 40]. We expanded on the range of thermal stability profiles for the native C-rich ILPR variants [25], with the two mutant sequences (Figure S3). The ILPR 1C_mut_ showed an increased thermal stability compared to the predominant ILPR 1C variant (Table S1, p <0.0001). However, ILPR 3C_mut_ has comparable thermal stability to ILPR 3C (Table 1, Figure S3, p=0.07).

Thermal difference spectroscopy (TDS) is commonly used to identify secondary DNA structures in solution, providing distinct spectral profiles that reflect on structural features. Both mutant ILPR sequences displayed TDS signatures with positive peaks at 240 and 265 nm and a negative peak at 295 nm (Figure S4), consistent with the presence of i-motif structures [41]. However, ILPR 6C presents as an intriguing candidate, exhibiting features of a hairpin in both CD spectra and TDS profile at pH 5.5, yet displaying a distinct melting profile in UV absorbance at 295 nm, an unusual characteristic for canonical hairpin structures. This suggests the presence of a minor population of non-canonical DNA conformations, such as i-motifs or triplexes, which are known to exhibit melting profiles at 295 nm due to base stacking interactions and structural transitions specific to these motifs [42].

The complementary native G-rich ILPR sequences have been previously studied [12, 43-47]. One study identified a correlation between G-quadruplex conformation and selective binding affinity of insulin and insulin-like growth factor [47]. We previously reported the biophysical characterisation of the G-rich ILPR sequences showing that only 3 of the native 11 ILPR sequences in the human insulin promotor element form stable G-quadruplexes, here renamed to 1G, 2G, and 3G [25]. We sought to determine whether the shifts in pH_T_ that were associated with mutations in loops of the 1C and 3C sequences were mirrored by changes in the complementary mutant G rich strand i.e. 1G_mut_ and ILPR 3G_mut_. We used QGRS mapper [48] to predict the stability of both mutant ILPR G-rich mutants, which exhibited a QGRS score of 63 as 1G, 2G, and 3G (Table 2).

**Table 2.**
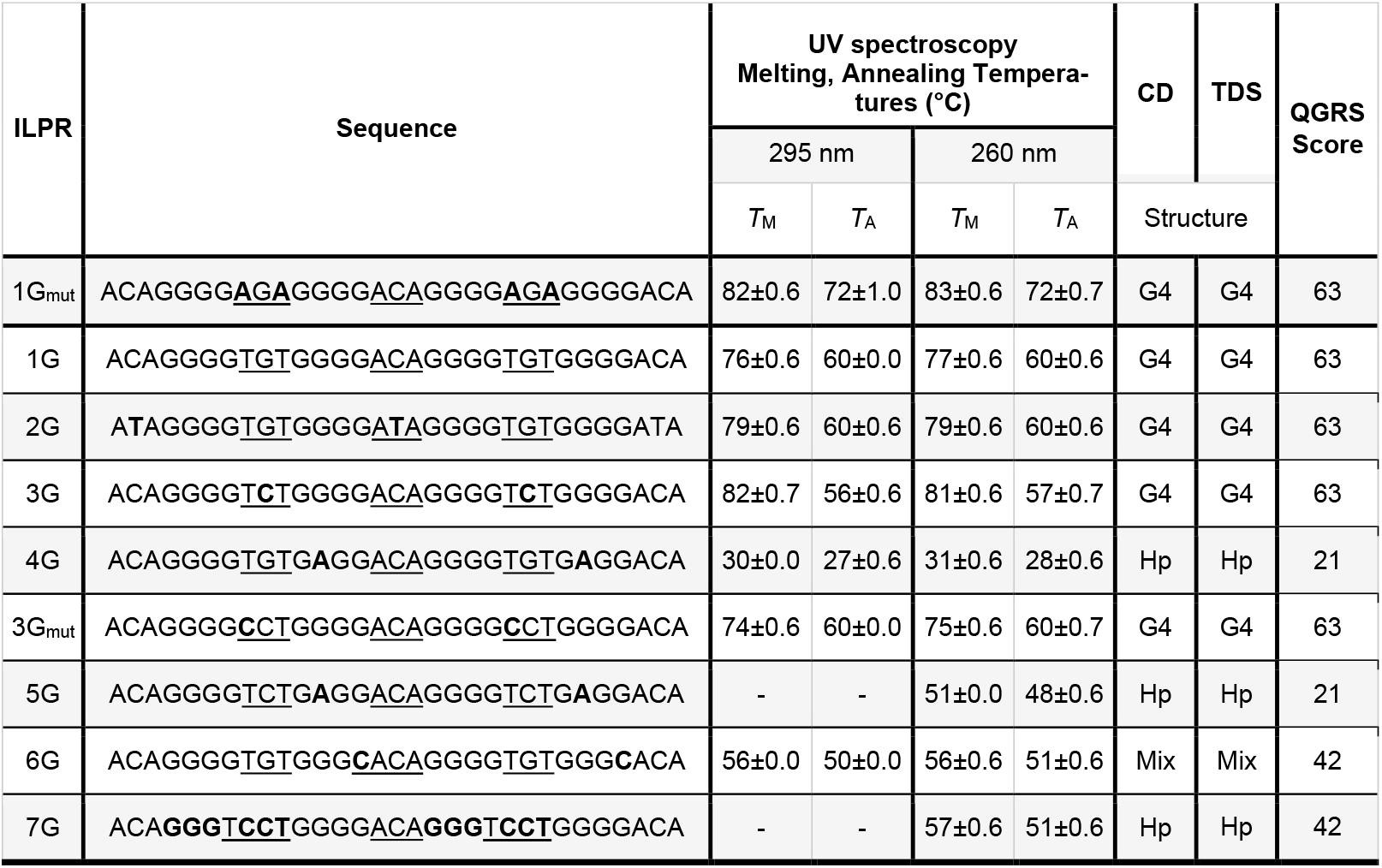
Biophysical characterisation via CD spectrum, UV melt and anneal, thermal difference spectrum, and QGRS mapper score of G-rich ILPR mutants in comparison with naturally occurring ILPR variants as published in Guneri et al., 2024. (− = not detected, GQRS Score = Quadruplex forming G-Rich Sequences Score, Melting Temperature = T_M_, Annealing Temperature = T_A_, Transitional pH = pHT, G-quadruplex =G4, Hairpin = Hp, mixture of G4 and Hp = Mix)

The G-rich ILPR sequence variants were characterised by circular dichroism (CD) spectroscopy in 10 mM NaCaco buffer (pH 7.0) containing 100 mM KCl, NaCl, or LiCl, to assess cation preferences, commonly used to observe cation dependent G-quadruplex topologies. Interestingly, the CD spectrum of ILPR 1C_mut_ has the typical profile for a parallel G-quadruplex with a positive peak at 264 nm and a negative peak around 240 nm (Figure S5A). However, ILPR C1 is a mixed population of parallel and antiparallel G-quadruplexes, presenting with a negative peak at 240 nm, and positive peaks at 263 nm and 295 nm Figure S5B). In potassium chloride, ILPR 3G_mut_ exhibits a mixed population with a higher tendency towards parallel G-quadruplex topology than antiparallel G-quadruplex topology, while in sodium chloride and lithium chloride the mixed topology has a higher preference for antiparallel G-quadruplex topology over parallel G-quadruplex topology (Figure S5E). These structural preferences in the three tested cations were reversed for the ILPR 3G variant. UV-melt analysis of 1G_mut_ shows a significantly increased melting and annealing temperature (Figure S6) compared to the native ILPR 1G variant (Table 2, p>0.001). ILPR 3G_mut_ has a significantly lower melting and anneal temperature compared to the native ILPR 3G variant (Table 2, p>0.001) [25]. The thermal difference spectrum provides profiles of typical Gquadruplex formation for both tested G-rich ILPR mutant sequences (Figure S7) [41].

### Both G-quadruplex and i-motif structures are required for reporter gene transcription

To further investigate the hypothesis that i-motifs and G-quadruplexes within the ILPR function as regulatory elements modulating insulin transcription, we selected ILPR variants predicted to adopt distinct combinations of DNA secondary structures: G-quadruplex and i-motif; hairpin and hairpin; and i-motif and hairpin. Despite examining over 100 ILPR-like sequences, we did not identify any combinations in which formation of G-quadruplex structure was accompanied by a hairpin in the C-rich complementary strand. We speculate that G-quadruplexes may be more prone to disruption by sequence mutations, which is also supported by previous studies showing that even single nucleotide substitutions within Grich regions—such as those in the c-MYC promoter—can destabilise G-quadruplex formation and shift the structural equilibrium toward alternative conformations, including hairpins [49-51]. The nine characterised ILPR sequences were cloned upstream of the human insulin promoter to regulate firefly luciferase expression in reporter gene assays, where luciferase activity serves as a proxy for promoter activation. Insulin-secreting β-cells respond to elevated glucose levels, enabling the assessment of ILPR-mediated regulation under physiologically relevant conditions linked to glucose homeostasis [52]. The rat insulinoma-derived cell line INS-1, widely used to study β-cell function, lacks an intrinsic ILPR or any known homologous sequence, making it a suitable model system for dissecting ILPR-mediated transcriptional regulation [25, 53, 54]. INS-1 cells were co-transfected with ILPR–luciferase constructs and a Renilla luciferase plasmid to normalise transfection efficiency. Following overnight serum and glucose starvation, the cells were exposed to either low (2.8 mM) or high (16.2 mM) glucose for four hours to assess glucose-responsive transcriptional activity, consistent with established protocols [25, 55-57]. We tested three categories of ILPR-regulated firefly reporter vectors: those forming G-quadruplexes and i-motifs (ILPR 1, 1_mut_, 2, 3 and 3_mut_), those forming hairpins on both DNA strands (ILPR 6 and 7), and those forming an imotif and a hairpin (ILPR 4 and 5).

Under low glucose conditions (2.8 mM), ILPR 1_mut_ showed a significantly increased firefly luciferase activity compared to ILPR 1 (Figure 2, p < 0.001). In contrast, reporter constructs driven by ILPRs 2, 3, 4, 3_mut_, 4 and 5 showed no significant differences in expression levels relative to ILPR 1 (p > 0.24), while ILPR 6 and 7 exhibited significantly reduced firefly activity (p < 0.001). At high glucose concentrations (16.2 mM), ILPR 1_mut_ again displayed significantly increased expression compared to ILPR 1 (p < 0.001). Meanwhile, ILPR 2 (p = 0.005), 3, and 3_mut_ demonstrated significantly decreased firefly activity (p < 0.001). No significant changes were observed for ILPR 4, 5, 6, or 7 under high glucose conditions. Interestingly, reporter constructs driven by ILPR sequences known to form i-motif and G-quadruplex structures *in vitro* (ILPR 1, 1_mut_, 2, 3, and 3_mut_) showed a significant overall increase in firefly luciferase activity when glucose levels were high (16.2 mM) compared to low glucose conditions (p < 0.001). The findings for ILPR 1, 2, 4, and 5 are consistent with our previous study [25] where we observed that sequences which form both G-quadruplexes and i-motifs in *in vitro*, showed increased firefly reporter gene expression in transfected INS-1 cells when treated with high glucose. In contrast, sequences, which form hairpins on both strands, showed no significant changes in expression between low and high glucose conditions.

**Figure 2.**
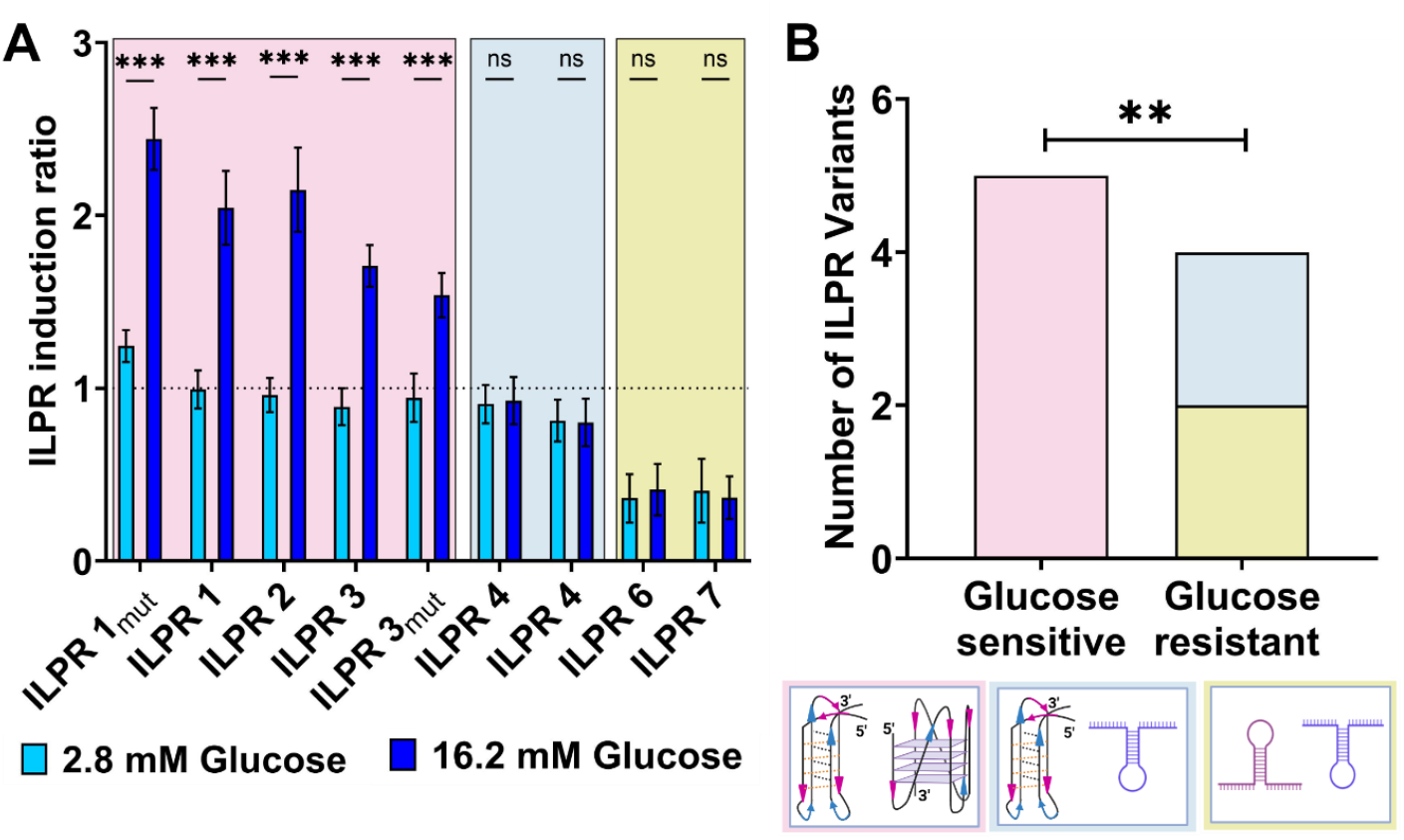
Dual Luciferase reporter gene assay in co-transfected INS-1 cells to determine response to 2.8 mM glucose (bright blue) and 16.2 mM glucose (dark blue) of firefly luciferase regulated by i-motif/G-quadruplex (pink background), hairpin/hairpin (blue background), and iM/hairpin (gold background) forming ILPR variants (A). Mean ± SEM (7 biological repeats each with 3 technical repeats); 2-way ANOVA with Holm-Šidák post Hoc corrections; p< 0.001***, ns > 0.1. Fisher’s exact test displays a significant association (p=0.008) between G4/iM formation and Glucose response in comparison to variants incapable to form both higher order non-canonical structures that remain non-responsive to glucose levels in cell-based reporter gene expression assay (B).

Plasmids containing ILPR 1, 1_mut_, 2, 3, and 3_mut_ showed significantly increased firefly gene expression in response to high glucose levels compared to low glucose conditions (p < 0.001). All five ILPR variants share underlying sequences capable of forming both i-motif and G-quadruplex structures and responded to glucose level changes in a similar manner. However, the increase in expression was significantly greater for ILPR 1_mut_, which also exhibited the most stable non-canonical DNA structures in the biophysical characterisation (p < 0.001). Importantly, reporter constructs containing ILPR sequences that do not form i-motif or G-quadruplex structures (ILPR 6 and 7) showed no changes in firefly gene expression under high glucose conditions (p > 0.99). Similarly, ILPR 4 and 5, which are associated with sequences which form weak i-motif and complementary G-rich hairpin structures *in vitro*, also showed no significant changes in response to glucose level variation (p > 0.99), indicative that this combination of structures does not trigger expression. These data highlight the potential significance of sequence variation within the ILPR, suggesting that the distinct DNA structures formed—such as i-motifs, G-quadruplexes, and hairpins—may influence the responsiveness to glucose. The findings suggest that even minor sequence alterations can result in substantial differences in both the formation and stability of DNA secondary structures, underlining the regulatory importance of both i-motif and G-quadruplex structures in controlling reporter gene expression.

Next, we interrogated whether glucose responsiveness was significantly associated with structural features of the ILPR sequences. Specifically, we compared glucose-sensitive and glucose-resistant ILPRs across three structural categories: (1) i-motif and G-quadruplex forming sequences, (2) hairpins on both strands, and (3) i-motif and G-rich hairpin combinations regulating firefly reporter gene expression. To assess the association between glucose responsiveness and the ability to form i-motif/G-quadruplex (imotif/G-quadruplex) structures, we applied a Fisher’s exact test (Figure 2B). Fisher’s exact test was chosen because it is particularly well-suited for small sample sizes and categorical data, such as the binary classification of ILPR variants into glucose-responsive vs. non-responsive, and i-motif/G-quadruplex-forming vs. non-forming groups. Unlike chi-square tests, Fisher’s exact test does not rely on large sample assumptions and provides an exact p-value, making it the most reliable option for detecting statistically significant associations in our dataset [58]. The analysis revealed a significant association between glucose responsiveness and the ability of ILPR sequences to form i-motif and G-quadruplex structures (p = 0.008).

### i-Motif stability modulates transcriptional activity

To better understand the contribution of secondary DNA structures to the regulation of gene expression, we aimed to determine whether i-motifs or G-quadruplexes play a more prominent role in modulating ILPR-driven promoter activity in response to glucose. We performed correlation analyses between biophysical parameters of structural stability and reporter gene expression under both low and high glucose conditions. Pearson’s correlation coefficient (r) was chosen as the statistical test to assess the linear relationships between, normally distributed variables [59]. In this case, biophysical measurements of imotif and G-quadruplex stability (*T*_M_) and quantitative firefly gene expression levels (Transcription induction ratio). This test allows us to evaluate whether increases in structural stability correlate with increased reporter activity across different ILPR variants. Pearson’s correlation analysis revealed a statistically significant positive correlation between i-motif stability, expressed as transitional pH (Figure S4), and ILPR-regulated firefly gene expression at both low (2.8 mM) and high (16.2 mM) glucose levels (Figure S8, p = 0.03; Figure 3B, p = 0.01). The melting temperature provides a direct comparison between both noncanonical DNA structures. The melting temperature is an additional parameter to express i-motif stability, a strong positive correlation was again observed with reporter gene expression under both low (Figure 3A, p < 0.001) and high glucose treatment (Figure 3B, p < 0.001). In contrast, no significant correlation was found between G-quadruplex stability, as measured by melting temperature and ILPR-induced gene expression at either low (Figure 3C, p = 0.31) or high glucose (Figure 3D, p = 0.24) levels. These findings suggest that i-motif stability is more closely associated with promoter responsiveness in this context, highlighting its potential regulatory relevance over G-quadruplexes in glucose-sensitive transcriptional control.

**Figure 3.**
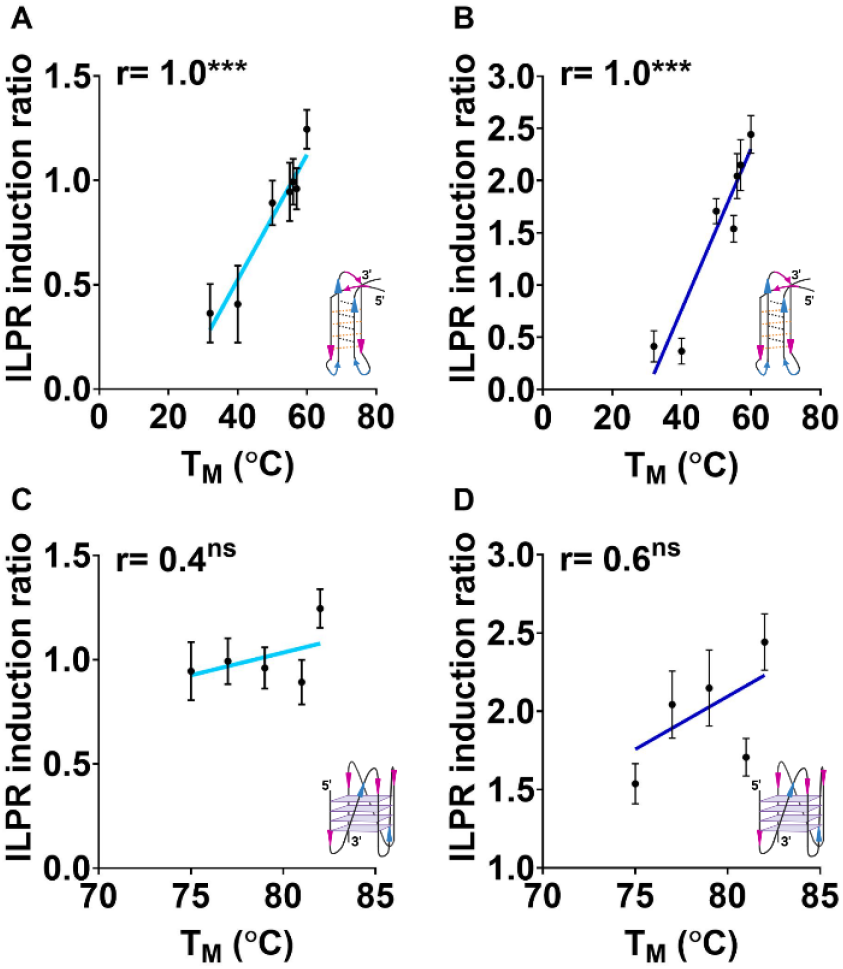
Pearson’s correlation between melting temperature C-rich ILPR variants capable of forming i-motifs and melting temperature of G-rich variants capable G-quadruplex folding and corresponding ILPR induction ratio in dual luciferase reporter gene assay in presence of 2.8 mM Glucose (A, C) and 16.2 mM Glucose (B, D). Data shown as Mean ± SD (n=6 for reporter gene assay, n=3 for biophysical data), student t-test; ns > 0.1, p< 0.001***.

Our findings support the idea that seemingly small sequence alterations within ILPR sequences can significantly impact the formation and stability of non-canonical DNA structures, thereby modulating glucose-responsive gene expression. Notably, ILPR 1_mut_ exhibited a stronger transcriptional response and greater structural stability of i-motif and G-quadruplex formations compared to wild-type ILPR 1, suggesting that even subtle sequence changes can shift the structural equilibrium toward conformations with enhanced regulatory capacity. This observation is consistent with previous studies showing that singlenucleotide variations or loop modifications in G-rich regions of the human telomeric, Bcl-2, and c-Myc promoter sequences can disrupt G-quadruplex formation and instead favour alternative structures such as hairpins, depending on sequence context and ionic conditions [39, 51, 60]. These findings reinforce our interpretation that the differences in glucose responsiveness observed between ILPR 1 and 1_mut_, as well as between ILPR 3 and 3_mut_, may be attributed not only to changes in structural stability but also to a potential shift toward parallel G-quadruplex conformations, which may be more transcriptionally active [61-63]. In contrast, variants such as ILPR 4, ILPR 5, ILPR 6, and ILPR 7, which predominantly adopted hairpin-like structures, showed minimal or no transcriptional activation in response to elevated glucose. Our data thus align with and extend prior reports by demonstrating that structural shifts, driven by minor sequence changes—can have a profound functional impact in the context of glucose regulation.

On the complementary C-rich strand, our data show that mutations in ILPR variants can alter i-motif stability, but i-motif formation is less frequently disrupted compared to the G-quadruplex. This suggests that i-motifs may possess greater sequence tolerance, maintaining folded structures even in the presence of point mutations. For instance, ILPR 1_mut_ exhibited an increased transitional pH and melting temperature compared to ILPR 1, indicating enhanced i-motif stability, which positively correlated with higher reporter gene expression under both low and high glucose conditions (Figures 3A–B). This supports the hypothesis that the structural integrity of i-motifs directly contributes to transcriptional responsiveness. Previous studies have similarly shown that single-base changes or loop bulges in i-motif-forming sequences can subtly modulate folding stability without fully abolishing structure formation. In the Bcl-2 and c-MYC promoters, for example, mutations created bulged loops or altered loop length and position, leading to changes in folding kinetics and pH responsiveness without completely preventing i-motif formation [39, 64]. Kaiser *et al*. investigated the KRAS promoter region and identified that the C-rich mid-region forms a dynamic equilibrium between i-motif, hairpin, and hybrid structures. Their study demonstrated that these structural forms are influenced by pH and that the i-motif structure can interact with the transcription factor hnRNP K to modulate KRAS transcription [65]. This suggests that structural variations, including bulges within the i-motif, can impact folding kinetics and transcriptional regulation. These findings align with our observation that ILPR variants forming stable i-motifs, such as ILPR 1_mut_, are more transcriptionally active than those forming weaker or unstable structures. Together, this reinforces the emerging role of i-motifs as tuneable, responsive DNA elements capable of regulating gene expression in a glucose-sensitive manner.

G-quadruplexes and i-motifs are non-canonical DNA structures that can form within genomic DNA, yet their presence is temporally regulated throughout the cell cycle, indicating distinct functional roles. Gquadruplexes tend to form during S-phase when DNA is unwound for replication, capable of stalling polymerases and influencing transcription and replication timing [22]. Conversely, i-motifs, which form on the complementary C-rich strand, are predominantly observed during late G1 to early S-phase [22], suggesting a more dynamic and regulatory role. The formation of the G-quadruplex and i-motif within the ILPR element has been shown to be mutually exclusive, likely due to steric hindrance [26, 66]. Nonetheless, these non-canonical structures may act in concert by providing binding sites for regulatory proteins, including transcription factors such as Pur-1 [8], as well as insulin and insulin-like growth factor 2 (IGF-2) [12]. The potential dynamic and context-dependent nature of G-quadruplex and i-motif formation suggests a potential regulatory mechanism finely tuned to the physiological state of the cell. This aligns with the observation that pancreatic β-cells possess a tightly controlled and protected cell cycle, with limited responsiveness to proliferative stimuli to preserve their differentiated state and insulin-secreting function [67]. These constraints reinforce the need for highly regulated transcriptional programs to maintain glucose homeostasis and support a model in which dynamic non-canonical DNA structures like those within the ILPR contribute to fine-tuned gene regulation in β-cells. Zeraati *et al*. demonstrated the presence of i-motifs in human nuclei with peak formation in late G1 phase, implying that i-motifs respond to finely tuned cellular cues, such as pH or redox state [22, 68]. Further evidence underscores this regulatory significance by showing that it is the stability of i-motifs, rather than G-quadruplexes, that correlates with spontaneous deletion events in human cells. These deletions were more frequent at loci with stable imotifs, suggesting that the persistence of i-motif structures may interfere with genomic processes such as repair or replication [69]. Collectively, these findings support a model in which G-quadruplexes serve as on/off switches, while i-motifs act as tuneable regulators, with the potential to impact genome stability and transcription.

## Conclusions

This study indicates that both i-motif and G-quadruplex structures are essential for transcriptional regulation. The current working hypothesis is that G-quadruplexes act as on/off switches, while i-motifs act as tuneable regulators. This work further emphasises the importance of considering G-quadruplex and i-motif DNA structures as one dynamic system and has implications in both the regulation of gene expression and the potential of i-motifs and G-quadruplexes as therapeutic targets.

## Supplementary Data

Circular dichroism, UV-melting, thermal difference spectra and Pearson’s correlation for transitional pH.

## Data Availability

Following publication, all data is available on Zenodo DOI:10.5281/zenodo.15320278.

