## Supplementary Information for "G-Quadruplexes Act as an On/Off Switch While i-Motifs Regulate Insulin Expression in Reporter Gene Assays"

### Figures

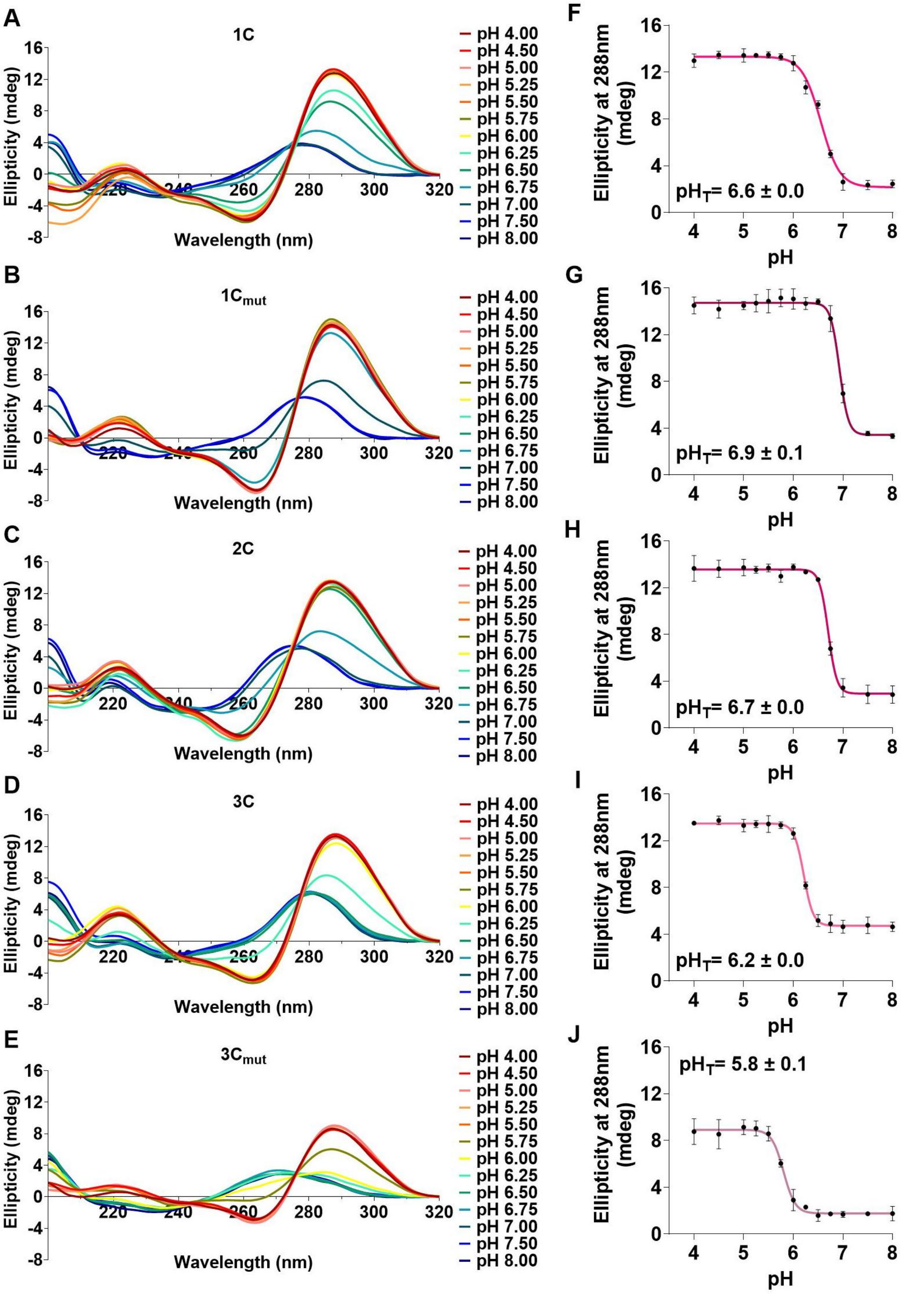


**Figure S1.** A-E) CD spectroscopy of C-rich ILPR sequences 1C, $1C_{mut}$, 2C, 3C, $3C_{mut}$. 10 μM DNA in 10 mM NaCaco 100 mM KCl and pH as indicated. F-J) Corresponding plot ellipticity of the experimental repeats (n=3, mean ± SD) at 288 nm at the measured pH conditions to determine transitional pH ($pH_T$) from the inflection point of the Boltzmann sigmoidal curve. Source data for this figure are provided as a Source Data file.

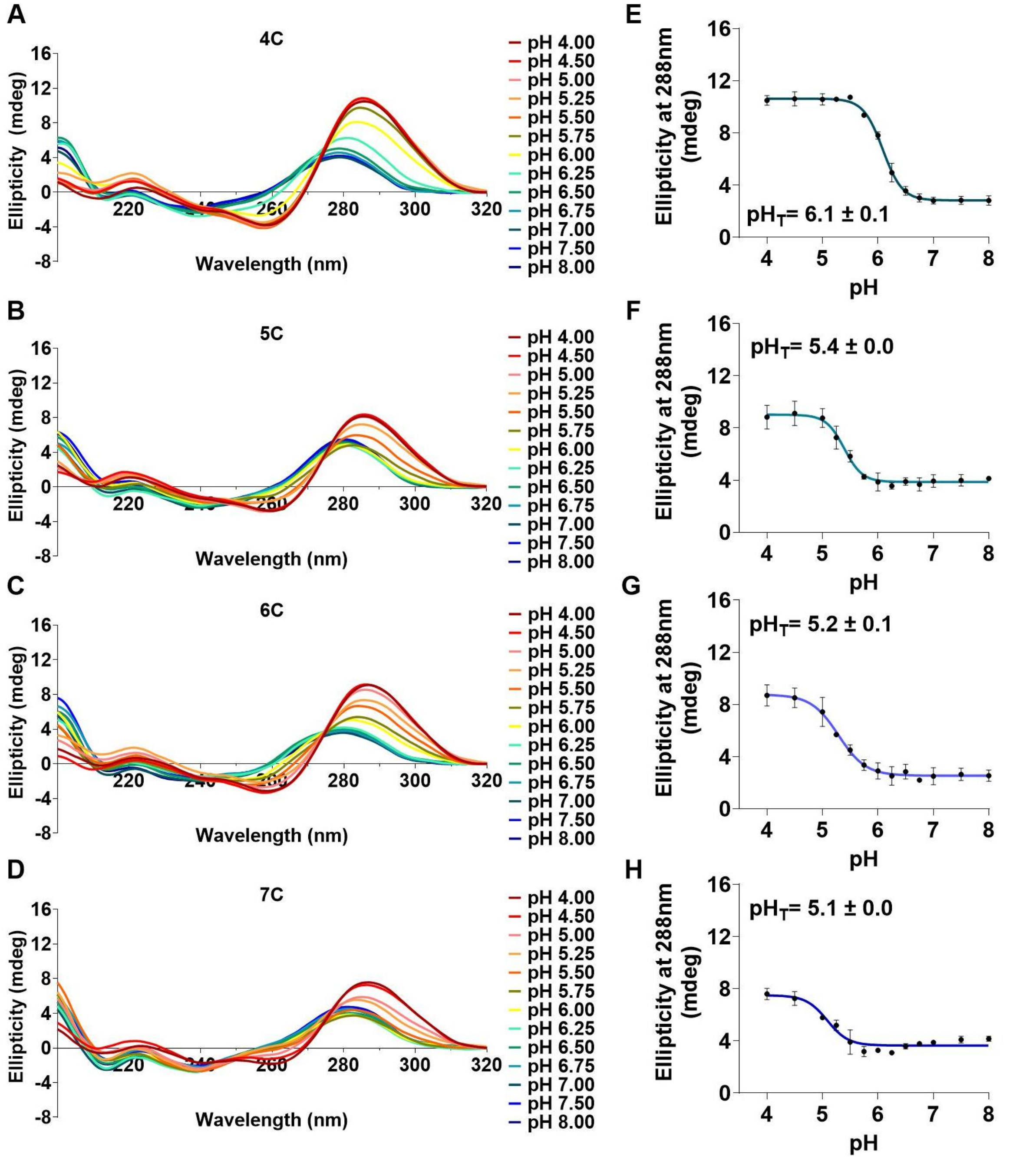


**Figure S2.** A-D) CD spectroscopy of C-rich ILPR sequences 4C, 5C, 6C, and 7C. 10 μM DNA in 10 mM NaCaco 100 mM KCl and pH as indicated. E-H) Corresponding plot ellipticity of the experimental repeats (n=3, mean ± SD) at 288 nm at the measured pH conditions to determine transitional pH ($pH_T$) from the inflection point of the Boltzmann sigmoidal curve. Source data for this figure are provided as a Source Data file.

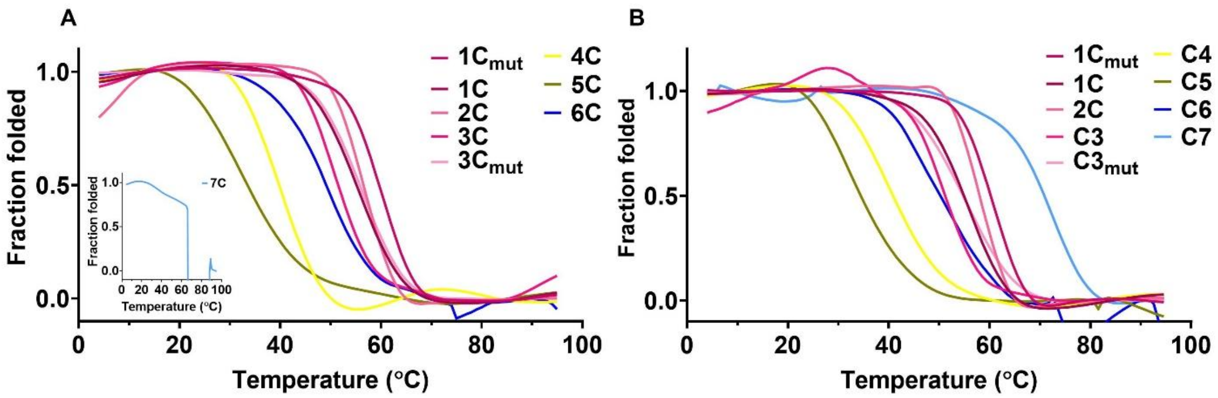


**Figure S3**. Fraction folded UV-melt recorded at 295 nm (A) and 260 nm (B) spectra analysis was performed using 2.5 μM DNA in 10 mM NaCaco 100 mM KCl at pH 5.5. The insert in A shows the lack of melting profile at 295 nm for 7C variants. Source data is available.

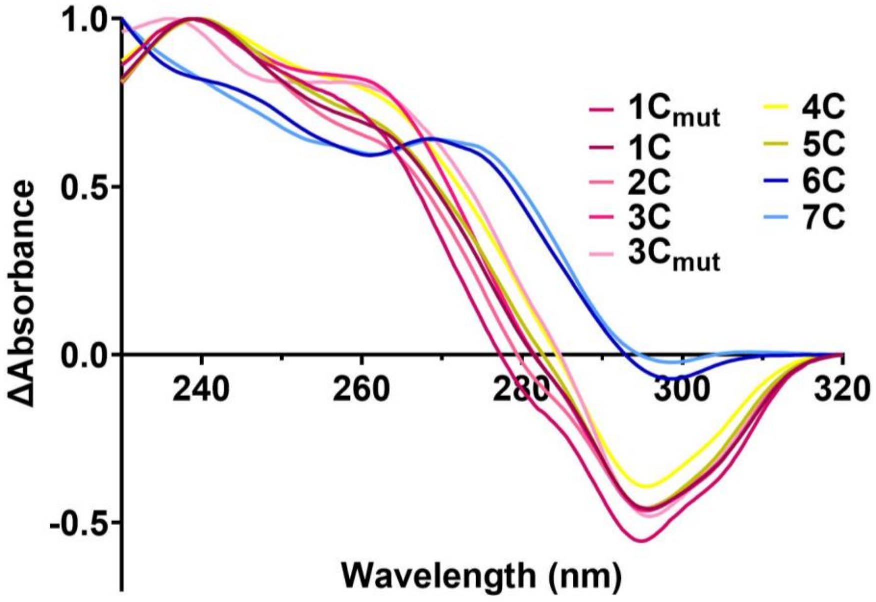


**Figure S4.** Thermal Difference Spectra of C-rich ILPR variants published in Guneri *et al*. for comparison to mutant ILPR C-rich sequences annealed as 2.5 μM DNA in 10 mM NaCaco 100 mM KCl at pH 5.5. Source data is available.

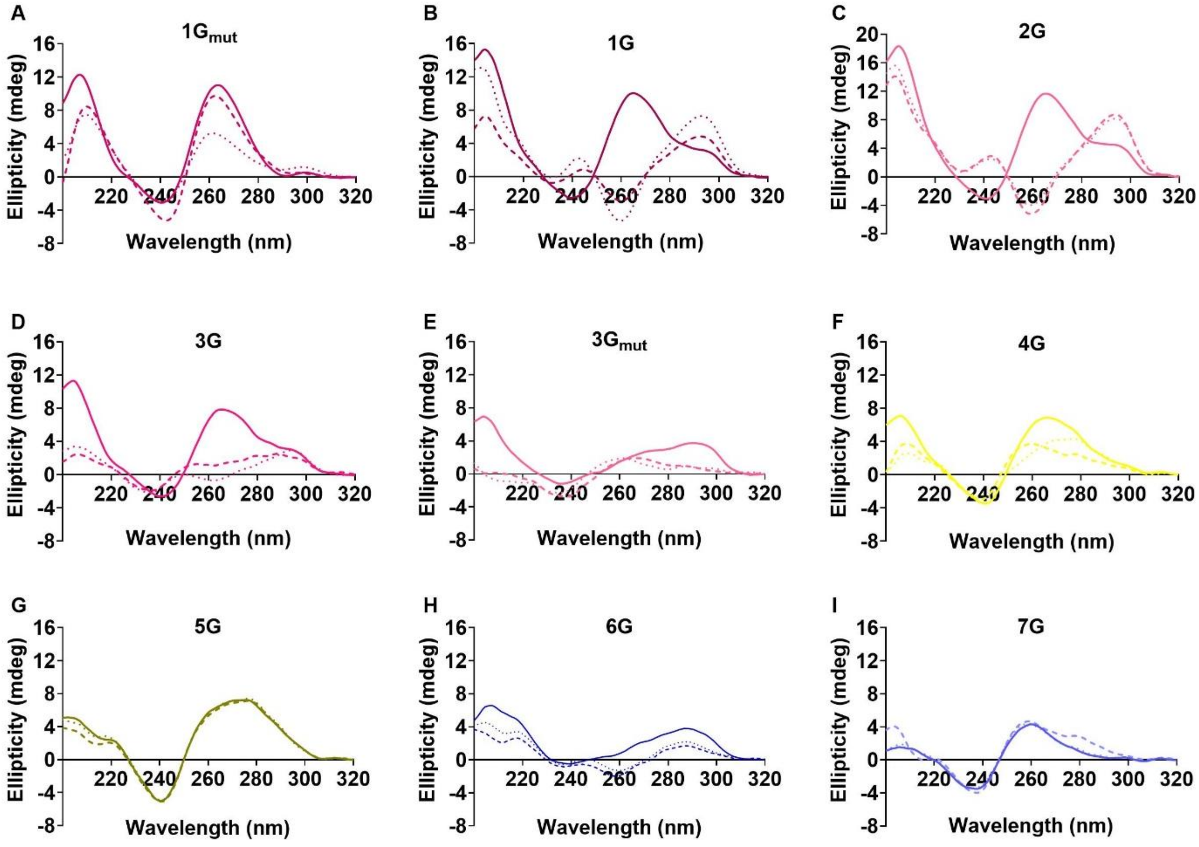


**Figure S5.** Biophysical characterisation of G-rich ILPR mutant sequences for non-canonical DNA structure formation and cation dependency. CD spectra analysis was performed using 10 µM DNA in 10 mM NaCaco 100 mM KCl at pH 7.0 (solid line), 100 mM NaCl (dashed line), or 100 mM LiCl (dotted line). Source data is available.

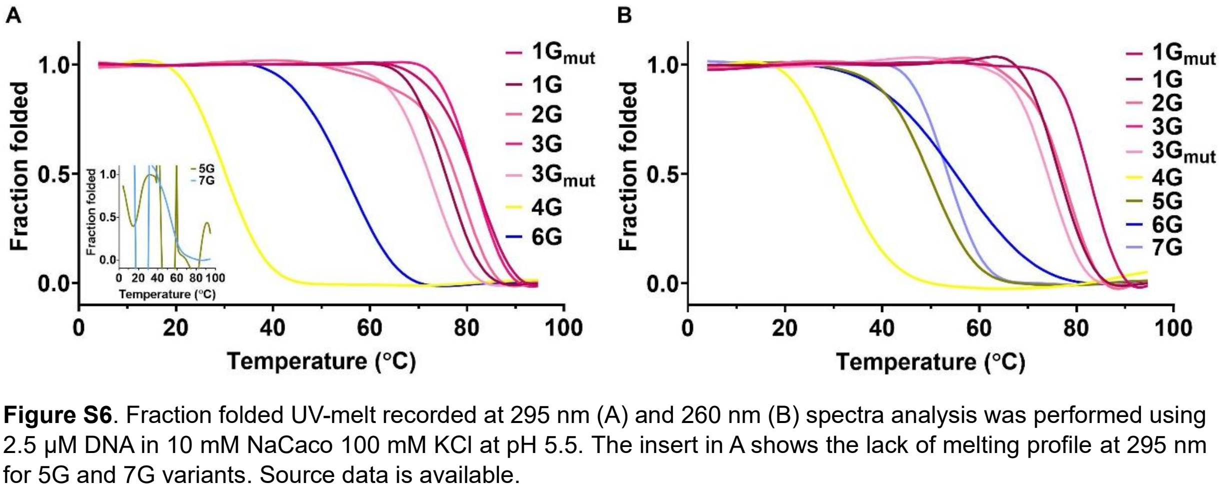


**Figure S6**. Fraction folded UV-melt recorded at 295 nm (A) and 260 nm (B) spectra analysis was performed using 2.5 µM DNA in 10 mM NaCaco 100 mM KCl at pH 5.5. The insert in A shows the lack of melting profile at 295 nm for 5G and 7G variants. Source data is available.

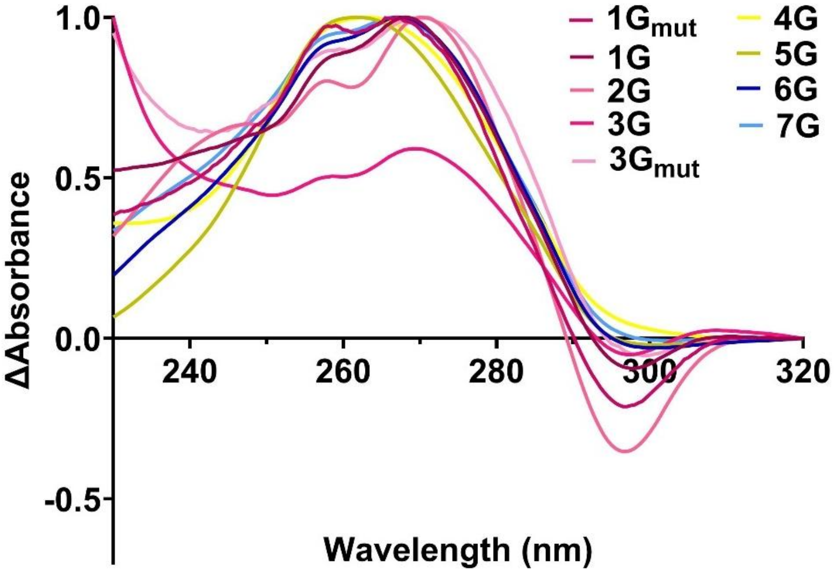


**Figure S7.** Thermal Difference Spectra of G-rich ILPR variants published in Guneri *et al*. for comparison to mutant ILPR G-rich sequences annealed as 2.5 μM DNA in 10 mM NaCaco 100 mM KCl at pH 5.5. Source data is available.

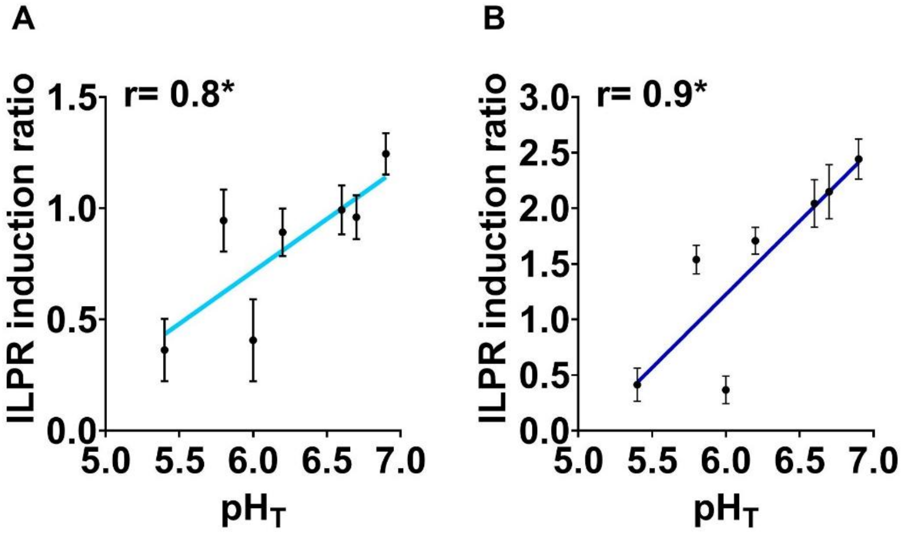


**Figure S8.** Pearson's correlation between transitional pH of C-rich ILPR variants capable of forming i-motifs and corresponding ILPR induction ratio in dual luciferase reporter gene assay in presence of 2.8 mM Glucose (A) and 16.2 mM Glucose (B). Data shown as Mean ± SD (n=6 for reporter gene assay, n=3 for biophysical data), student t-test , ns > 0.1, p>0.033*, p> 0.002**, p< 0.001***.
